# Serum proteomic approach for differentiation of frail and non-frail elderly

**DOI:** 10.1101/600262

**Authors:** Vertica Agnihotri, Abhishek Gupta, Rashmita Pradhan, G Venugopalan, Sailesh Bajpai, A. B. Dey, Sharmistha Dey

## Abstract

Frail elderly is very common in Indian society and extremely difficult to manage in the clinical practice. Blood proteomic study may able to improve the specific diagnostic profiles and therapeutic templates for improving quality of life in elderly. The purpose of present study is to differentiate between frail and non-frail elderly on the basis of serum markers. The proteomic profile of 10 frail and 10 non-frail elderly diagnosed according to deficit accumulation model of Rockwood was identified by 2D-electrophoresis and analyzed using ImageMaster 2DPlatinum7.0 software. The proteins were identified by MALDI-TOF/TOF and performed gene ontology study by PANTHER 7.0 software. Overall 105 spots were identified in the study groups. In frail 22spots and in non-frail 12spots were found to be differentially expressed. Mass spectroscopy analysis of 13 spots showed up-regulated Haptoglobin, Serum amyloidA1, TRAK1, sp110, NLRC3, MMP12, Mortalin, NDK3 and downregulated PSRC1, NKG2A proteins in frail elderly. The differential expression of proteins in frailty are mostly associated with pro-inflammation and is well-known that frailty syndrome increases all inflammatory parameters. It can be summarized from the present proteomic study that these proteins can be used as potential biomarkers for early detection of frailty in the elderly.

## 1. Introduction

The frailty syndrome has recently paying attention to the researchers and public health organizations as precursor and for the improvement of age-related conditions in older persons. Although physical frailty has shown to be related in epidemiological study, their pathophysiological mechanisms are not clear. The physical frailty is defined as “a medical syndrome with multiple causes and contributors that is characterized by diminished strength, endurance and reduced physiologic function that increases an individual’s vulnerability for developing increased dependency and/or death” [1]. Frailty is a state of global condition of impaired strength with increased susceptibility to various stressors. It upsurges the risk of hospitalization, falls and mortality. It makes a person dependent on others for maintaining their activities of daily living (ADL). There are two well described models of Frailty: Phenotype model given by Fried [2] which describes physical frailty and Cumulative deficit model given by Rockwood [3]. Clinical identification of frailty by any standard scales is possible only after considerable damage has been done at the cellular level. Non availability of any promising intervention till date is due to late diagnosis, as a result, intervention’s efficacy is poor. This is the vital reason behind the search for biomarkers to identify frailty at a much earlier stage, even before the clinical manifestations set in. This will also help in identifying the critical pathophysiological pathways which can be targeted by drugs. Before appearance of clinical phenotype multiple changes initiated at the cellular level. These cellular changes lead to changes in the normal expression of the molecules. These molecular changes in the tissues are reflected in the blood levels, which can be used to identify, predict and monitor frailty. Blood based biomarkers, tissue/cell based biomarkers and genetic biomarkers are being studied to find a consistent association with frailty in various population groups. Thus, in the next few years, a panel of biochemical or genetic markers may be used to predict and identify frailty in its earliest phase when it is amenable to some intervention.

The use of 2D proteomic approach for studying blood markers is in great demand. It is an exceptional and noninvasive method which has an ability to separate thousands of proteins straight away and detect even minor changes in expression profile of proteome in comparative groups which is necessary for early diagnosis of several diseases [4]. The application of proteomic studies has been reported to mark proteins associated with clinical pathologies that would further lead the way to biomarkers’ discovery.

Aging and frailty are often seen as pro-inflammatory state. Indeed today, aberrant inflammation is considered both an emerging hallmark of frailty and an enabling characteristic of aging. This has led to initiate works with inflammatory markers. Various studies have shown its association with haemoglobin, albumin, white blood count, cholesterol level, fibrinogen, transthyretin, retinol binding protein, tumour necrosis factor-alpha (TNF-α), Interleukin-6 (IL-6), C-22 reactive protein (CRP), Insulin like Growth Factor-1 (IGF-1), testosterone level, etc. [5-8]. Consistency of this association in a large cohort has to be done before making any final conclusion. Moreover these markers are highly non-specific and found in various other common degenerative conditions of old age. So, the research for a specific marker of frailty is wide open.

The application of genomics, proteomics and even metabolomics to the research on aging would increase our understanding of the origin of the different processes that contribute to the development of frailty in aging. Blood proteomic studies may be able to associate with disease process, thereby enabling the establishment of specific diagnostic profiles and therapeutic templates that could aid in improving quality of life in elderly.

## 2. Material and Methods

### 2.1. Study Group

Blood samples were collected from 20 participants aged above 60 years visiting Department of Geriatric Medicine, All India Institute of Medical Sciences, New Delhi. All participants were recruited with written informed consent and ethical approval was obtained by the Institute Ethics Committee (IESC/T-35/03.01.2014). All participants underwent comprehensive geriatric assessment (CGA) according to the “deficit accumulation model of Rockwood” and frailty status was determined by Frailty Index (FI) [9]. FI is derived from 36 item questionnaire assessing physical function, medical conditions, symptoms and functional impairments. Positive response to a question denotes ‘deficit’ in particular domain and this ‘accumulation of deficits’ is a well validated model used to determine frailty status in a wide range of population. Each positive response was scored with one point and FI is calculated as sum of number of deficits divided by total number of variables (n/36). A score of ≥ 0.25 (9 or more deficits) was used to identify frail subjects and the remainder was categorized as non-frail.

### 2.2. Serum and protein preparation

Two ml of blood sample was allowed to clot for 30min in microcentrifuge tube. Clotted samples were centrifuged at 3000 rpm for 10min and buffy coat was removed after that. Major interfering proteins were removed from the serum by the Multiple Affinity Removal Spin Cartridge HSA/IgG (Agilent Technologies, USA) as described into manual. Cold acetone (4 volumes) were added to sample (1 volume) for the precipitation of protein. Precipitated protein was dissolved into the rehydration buffer (8 M Urea, 2 M Thiourea and 4% CHAPS). Protein quantification was done by Bradford method (Bio Basic Inc. USA) using Bovine Serum Albumin (BSA) as a standard.

### 2.3. 2-Dimensional Polyacrylamide Gel Electrophoresis or 2 DE

*First Dimension***-** The sample was prepared by adding immobilized pH gradient (IPG) buffer (1.5 µl, pH 3-10), 0.25% Bromophenol blue dye (2 µl) and pinch of DTT into 300µg of protein containing rehydration buffer and made the final volume upto 250 µl with rehydration buffer. The sample was loaded on dry IPG (13 cm) strips (pH 3-10) for 15h at room temperature. After rehydration, iso-electric focusing (IEF) was performed on Ettan IPGphor3 system (*GE Healthcare*, Sweden) using the following program: 50 V (2h), 100V (1h), 500V (2h), 1000V (2h), 5000V (3h), 6600V (3h). In the next step, IPG strips were equilibrated by SDS buffer having 10 mg/ml DTT for 15min followed by second step equilibration with SDS buffer containing 25 mg/ml iodoacetamide for 15min.

*Second dimension*-SDS PAGE (12.5%) was run at 80 V until Bromophenol blue dye comes out of the gel. Colloidal Coomassie Brilliant Blue (CBB) G-250 staining was used to visualize the protein. Gel was fixed in ethanol and acetic acid solution in 40:10 ratio with final volume made upto 100ml with double distilled water and left the gel in solution for next 1h. Gel was stained overnight with colloidal CBB stain (containing 0.12% CBB G-250, 10% ammonium sulphate, 10% phosphoric acid, 25% methanol freshly prepared). Gel was scanned and analyzed using ImageMaster 2D Platinum 7.0 software (Agilent technologies, USA).

### 2.4. Image Analysis and Protein Identification by MALDI-TOF/MS

The spots for identification were excised based on a criterion of one fold change in normalized intensity from controls. The selected protein spot were identified by MALDI-TOF/TOF MS (Bruker Daltonics ULTRAFLEX III) by the Sandor Life Sciences Pvt. Ltd. Hyderabad. In brief, the gel slice was diced to small pieces and placed in new Eppendorf tubes. The gel pieces were destained using destaining solution for 3-4 times (10min intervals) until the gel pieces become translucent white. The gels were dehydrated using acetonitrile and Speedvac till complete dryness. The gel pieces were rehydrated with DTT and incubated for 1h. After incubation the DTT solution was removed. The gel pieces were now incubated with iodoacetamide for 45min. The supernatant was removed and the gel was incubated with ammonium bicarbonate solution for 10min. The supernatant was removed and the gel was dehydrated with acetonitrile for 10min and Speedvac till complete dryness. Trypsin solution was added and incubated overnight at 37°C. The digest solution was transferred to fresh eppendorf tubes. The gel pieces were extracted thrice with extraction buffer and the supernatant was collected each time into the eppendorf above and then Speedvac till complete dryness. The dried pepmix was suspended in TA buffer. The peptides obtained were mixed with HCCA matrix in 1:1 ratio and the resulting 2 ul was spotted onto the MALDI plate. After air drying the sample, it was analyzed on the MALDI TOF/TOF ULTRAFLEX III instrument and further analysis was done with FLEX ANALYSIS SOFTWARE for obtaining the PEPTIDE MASS FINGERPRINT. The masses obtained in the peptide mass fingerprint were submitted for Mascot search in “CONCERNED” database for identification of the protein. MASCOT takes into account the quality of the peaks submitted while performing the search and Proteins with a p-value less than 0.05 (statistical significance) are only given as Positive results. The Mass tolerance is used to control the false positive rate during analysis.

### 2.5. Bioinformatics Analysis

Relative spot intensities by comparison group were estimated using ImageMaster 2D platinum 7.0 software (GE Healthcare). Those showing more than or equal to one fold difference between groups (frail vs nonfrail) were selected and identified using MALDI-TOF. The functional analysis of identified proteins (13 proteins) was done on Gene Ontology (GO) using PANTHER 7.0 bioinformatics software platform. GO analysis produced information about biological, cellular and molecular functions of different proteins. For interactive studies, following proteins were analysed by means of STRING software.

### 2.6. Statistical Analysis

Statistical analysis was executed using Graph Pad version 5.0. The Fisher’s exact test was used for categorical variables and unpaired t-test for comparison of continuous variables. p-value ≤0.05 was considered statistically significant.

## 3. Results

### 3.1. Baseline data of the subjects

The characteristics of 10 non-frail and 10 frail subjects are presented in Table 1. A Rockwood criterion was used for clinical assessment and diagnosis of the study participants. Mean age of the non-frail was 77.25±6.01 (Mean±SD) years while mean age of frail was 81.75±6.54 (Mean±SD) years. The gait speed and grip strength were significantly lower in frail as compared to non-frail elderly.

**Table 1:** Demographic characteristics of Frail and Non-Frail.

| **Variable** | **Frail** | **Non-Frail** | **p Value\*** |
| --- | --- | --- | --- |
| **N** | 10 | 10 |  |
| **Age (Mean±SD)** | 81.75±6.54 | 77.25±6.01 | 0.174 |
| **Male** | 5 | 7 |  |
| **Female** | 5 | 3 | 0.409 |
| **BMI (Kg/m<sup>2</sup>) (Mean±SD)** | 20.17±2.01 | 22.50±1.88 | 0.015* |
| **OA Knee** |  |  |  |
| Yes | 6 | 0 |  |
| No | 4 | 10 | 0.010* |
| **Joint Pain** |  |  |  |
| Yes | 7 | 3 |  |
| No | 3 | 7 | 0.037* |
| **Fall** |  |  |  |
| Yes | 7 | 1 |  |
| No | 3 | 9 | 0.010* |
| **ADL** | 18.60±0.96 | 19.90±0.32 | 0.0008* |
| **Malnutrition Status** |  |  |  |
| Yes | 8 | 4 |  |
| No | 2 | 6 | 0.094 |
| **Gait Speed (m/s) (Mean±SD)** | 0.46±0.14 | 0.77±0.15 | 0.0002* |
| **Grip Strength (kg) (Mean±SD)** | 8.30±10.09 | 27.10±12.84 | 0.0019* |

Abbreviation: ADL = activities of daily living; BMI = body mass index; OA knee = osteoarthritis knee; \*p-value <0.05 is significant

### 3.2. 2-Dimensional Gel Electrophoresis

The serum samples were processed to remove major interfering protein present in the serum by Multiple Affinity Removal Spin Cartridge HSA/IgG before 2-DE. Total 105 spots were obtained through 12.5% acrylamide gel among them 13 spots were considered for mass spectroscopic analysis (Figure 1).

**Fig. 1.**
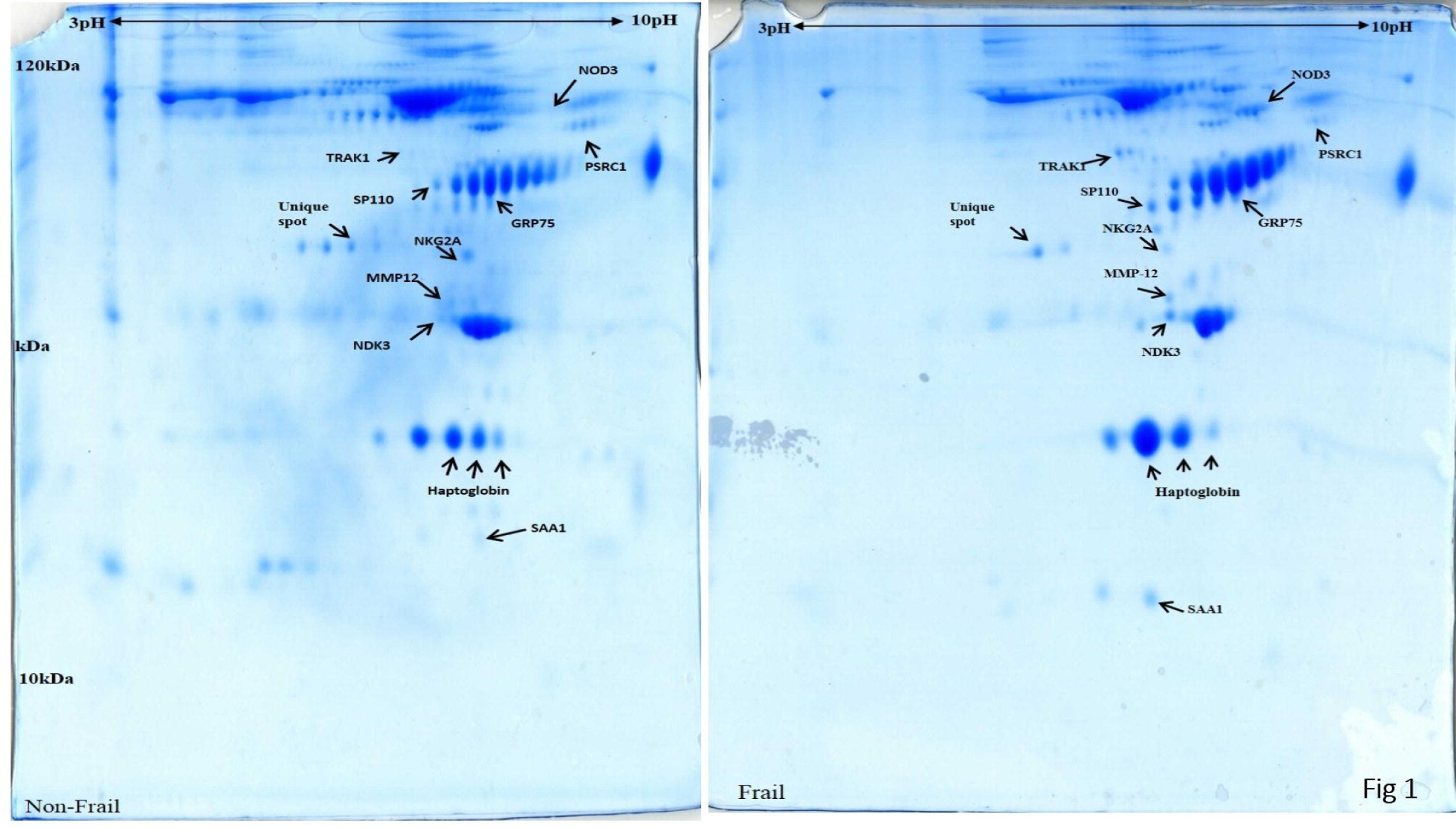
12.5% SDS gel image of Non-Frail and Frail serum samples: a) Non-Frail b) Frail

### 3.3 Protein Identification

One hundred five spots were seen on the Coomassie Brilliant Blue G-250 stained in non-frail and frail 2D gels. A number of differentially expressed and unique spots were seen on the gels. In frail, 10 overexpressed and 2 low expressed proteins were spotted as compared to non-frail gel and 1 unique spot was highly expressed in non-frail compared to frail but it was not found in the uniprot database. Spots were identified by MALDI-TOF/TOF/MS analysis. The list of the protein identified by the mass spectroscopy is shown in Table 2.The experiment was performed thrice.

**Table 2:**
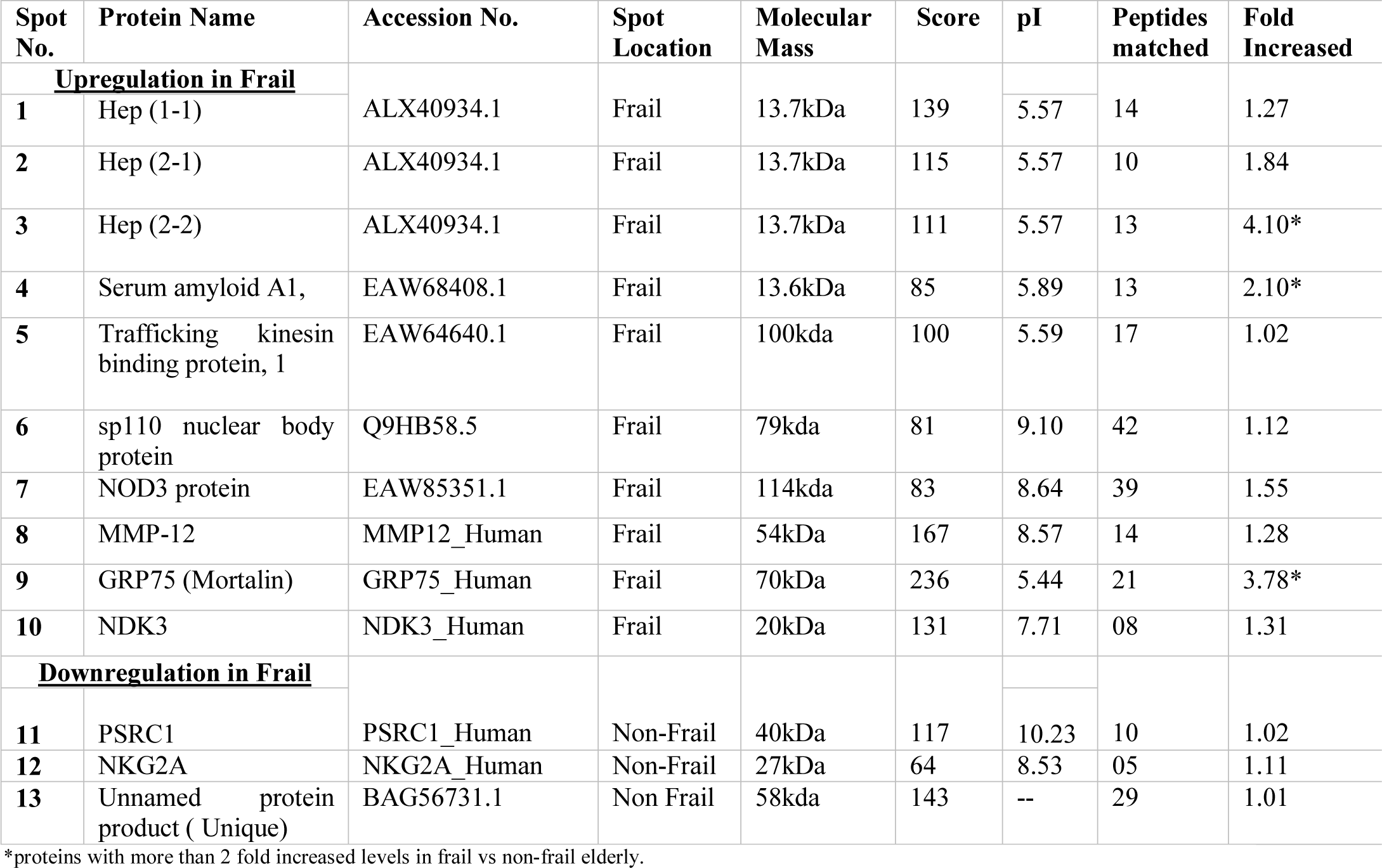
List of identified proteins from frail and nonfrail serum samples.

### 3.4. Functional Characterization of proteins

GO analysis, performed using PANTHER 7.0 bioinformatics software platform, had provided information about biological, cellular and molecular functions. The classification based on cellular component (Figure 3) showed that majority of the proteins are existing at cell projection part (37.5%), followed by extracellular region (25%), membrane bound organelle (25%) and at surface of membrane (12.5%). Classification for biological function bar graph depicted that maximum of the proteins are involved in mechanisms of stimulus function (23.1%), followed by biological regulation (15.4%), locomotion, metabolic process, cellular component organization (7.7%). The classification of molecular function showed that majority of proteins has catalytic activity (50%), binding (37.5%) and molecular transducer activity (12.5%). Inspite of sharing biological and molecular functions, STRING analysis did not give any favourable interaction among these proteins (data not shown).

**Fig. 2.**
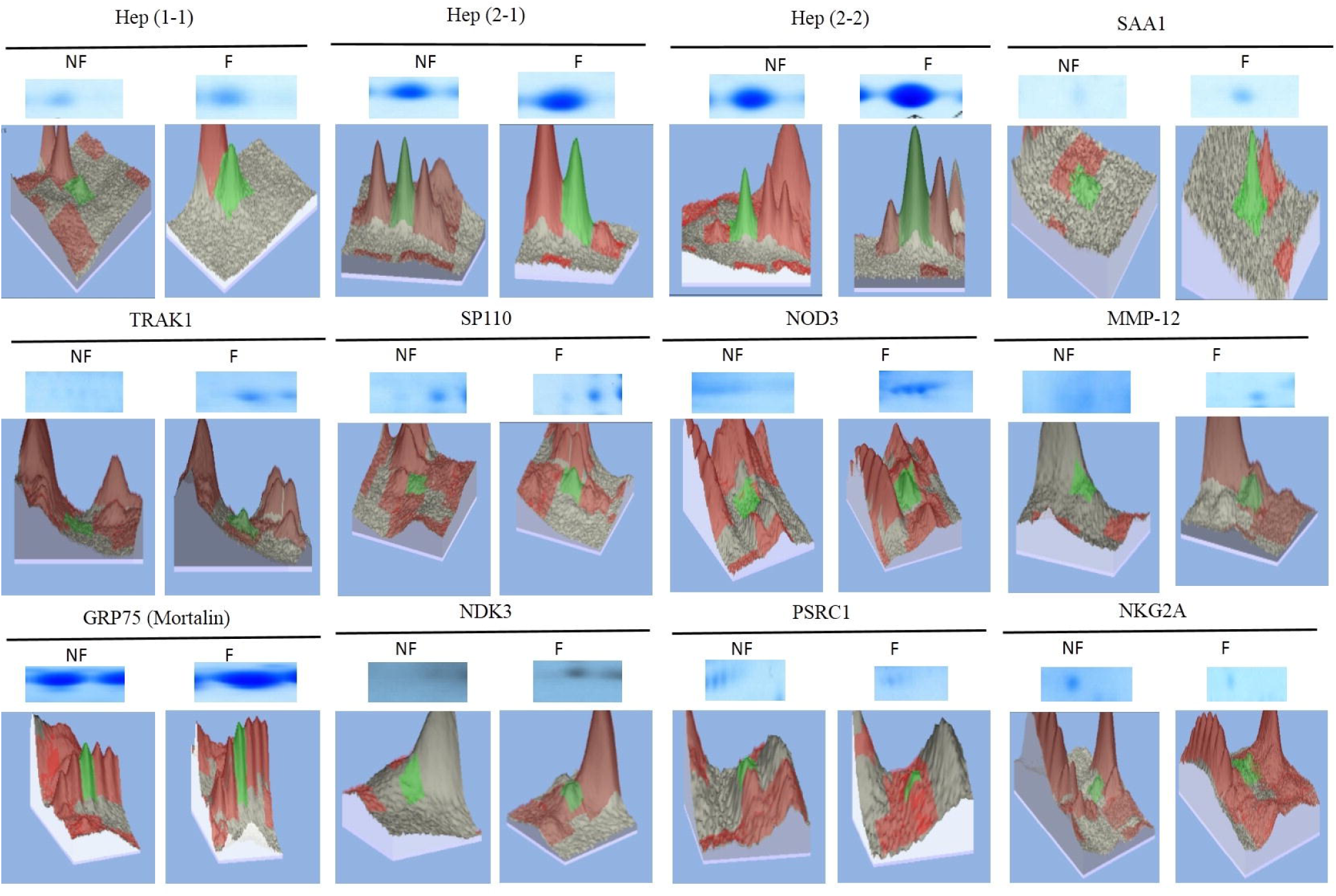
Upregulated and downregulated protein spots of pH3-10NL. Pair spots of each protein of Non-Frail and Frail were shown; 3D representation of spot volumes shown here was derived from Imagemaster 2D software; NF= Non-Frail, F= Frail.

**Fig. 3.**
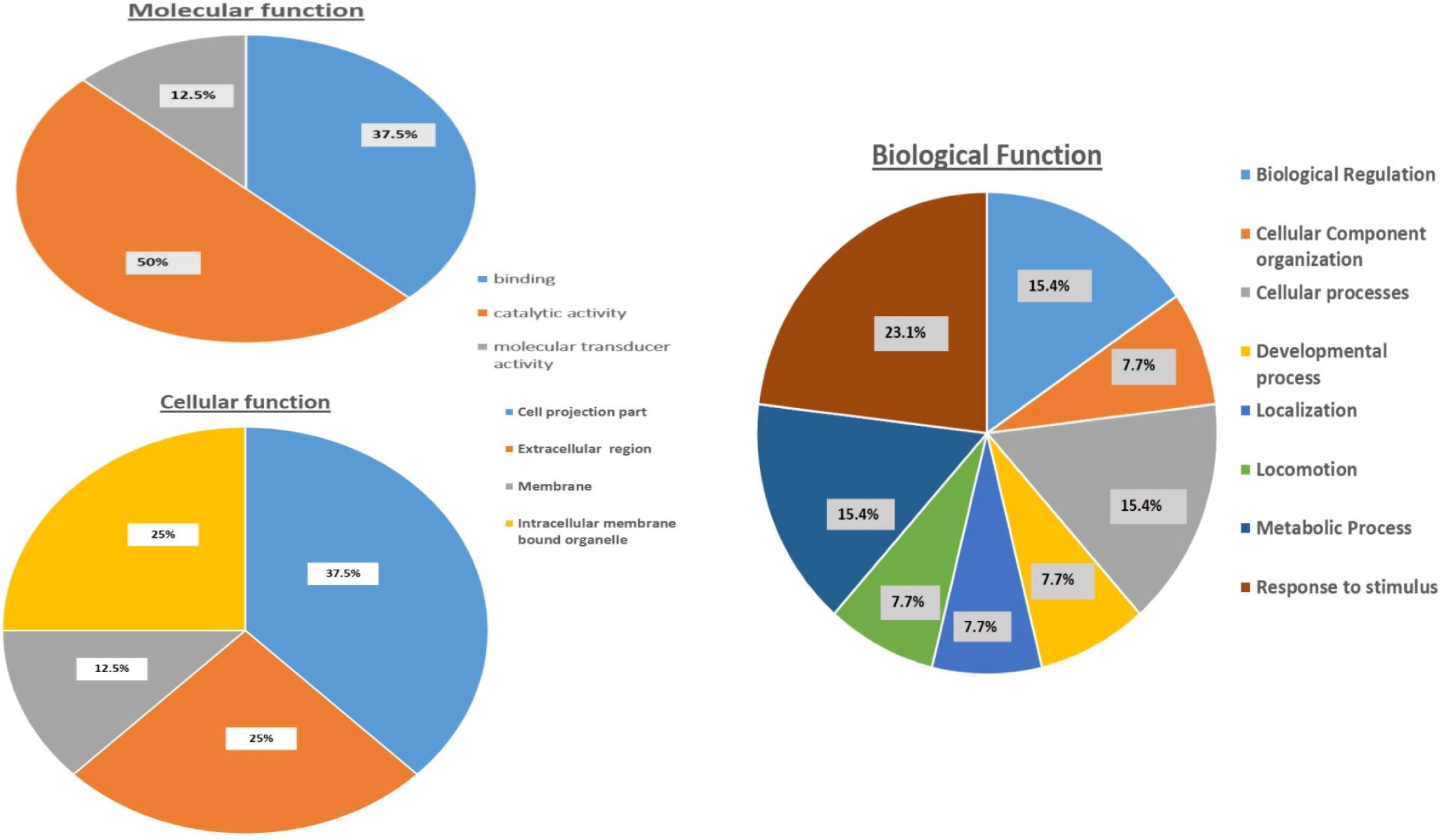
Gene ontology classification of proteins on the basis of their involvement in a cellular component, molecular function, and biological process using PANTHER 7.0 software.

## 4. Discussion

Frailty is an important and complex geriatric syndrome linked with inflammation. The connection between inflammation and frailty is complex as both increases linearly with advancement of old age. Frailty symptom increases all inflammatory parameters [10,11]. The inflammation in frailty affect muscular mass, increase BMI, reduce innate immune system, affect T-cell activity and increase mitochondrial activity which increases serum inflammatory level. The role of inflammation in the pathogenesis of frailty has not come to a conclusion yet. This study compared the serum protein profile of frail and non-frail elderly subject focusing on relation between inflammation and frailty. Differential expression of 34 proteins was found between frail and non-frail subjects.

In frail 18 spots were found which were not present in non-frail, likewise 53 spots were absent in frail. Thirteen protein spots were analyzed by MALDI-TOF/MS but one protein spot was not found in the uniprot database. Therefore, 12 spots were identified on the basis of considerable differential expressions present in both groups.

Among the 12 identified spots, 3 were haptoglobin protein isoforms which were found to be 1.27, 1.84, 4.10 fold respectively, overexpressed in frail compared to non-frail elderly. Three spots were identified as haptoglobin are the phenotypes of it; Hep (1-1), Hep (2-1) and Hep (2-2). Haptoglobin binds with free haemoglobin (Hb) in blood plasma and prevents the oxidative activity of Hb. Anemia is most frequently associated with frailty and there is a chemistry between inflammation and anemia in frailty. Haptoglobin is an antioxidant protein. It increases with inflammation and also in case of high level of ROS. While in previous report, differential expression of haptoglobin was not found between frail and non-frail by ELISA [12]. However, haptoglobin was found to be elevated in serum of aged rat [13].

Present study reports that Serum amyloid A (SAA) protein levels were 2.10 fold higher in frail as compared to non-frail samples. It secreted at the acute phase of inflammation by recruiting immune cells at inflammation site which support the previous study on serum. In a study, it is found that the elevated levels of SAA is the result of acute phase of inflammation [14], which eventually leads to decline in functionality of elderly, a symptom of frailty syndrome.

Mitochondria have been proposed to play a key role in aging in previous studies [15]. Trafficking kinesin binding protein (TRAK) family comprises many adaptors of kinesin which are responsible for neurons trafficking of the mitochondria. It regulates the mobility of mitochondria [16]. A recent immunocytochemical study reported that TRAK1 was widely localized in axons and has possibly distinct roles in mitochondrial transport in different neuronal subcellular compartments [17]. Several acute and chronic inflammatory diseases have always been linked to transformed mitochondrial function [18]. Our findings for the first time report that TRAK1 protein was found to be 1.02 fold overexpressed in frail compared to non-frail. It can be suggested that aberrant expression of TRAK1 in frail can lead to mitochondrial trafficking dysfunction in neurons and increase inflammation.

In frailty, significant alteration of innate immune system persist which upregulates the stress responsive inflammatory pathway genes. Although the underlying mechanism for frailty is likely multi-systemic, the activation of inflammatory pathways is likely to play a central role in its development and its related weakness to adverse health outcomes [19]. Sp110 is a member of the nuclear body (NB) components that functions as a nuclear hormone receptor transcriptional co-activator [20]. According to a study, Sp110 is induced by type I and type II interferons (IFNs). It shows an important role in IFN response and virus replication [21]. Our study found SP110 protein at higher level in frail compared to non-frail. This protein regulates gene transcription and plays a role in innate immune system.

Production of inflammatory cytokines is necessary for initiating innate immunity response. NLRC3 is the third family member of sub-group of NLRs family that negatively regulate inflammation [22]. It functions as an attenuating checkpoint for the production of inflammatory cytokines in macrophages [23]. NLRC3 diminished the production of inflammatory cytokines in vivo by attenuating NF-κB pathway. NLRC3 was originally identified as a negative regulator of T cell function. In this study, we found higher NLRC3 protein serum levels in frail comparing it with non-frail subjects. It could be suggested the high expression of this protein leads to aberrant behavior of innate immune response.

PSRC1, also known as DDA3, is directly regulated by p53 and has been reported in multiple processes in mitosis [24]. In a microarray experiment, PSRC1 was reported upregulated in human malignant peripheral nerve sheath tumor cells [25]. Higher expression of PSRC1 was reportedly downregulated expression of pro-inflammatory cytokines in RAW264.7 macrophage. PSRC1 activates β-catenin pathway and inactivates activity of nuclear transcription factor (NF-κB), which acts as a key gene in the regulation of inflammation. As reported in an in-vivo study, PSRC1 overexpression decreased the plasma levels of TC, TG, LDL-C, TNF-α, IL-1β and IL-6, increased the plasma HDL-C levels and improved HDL function [26]. Consistent with above studies, our results show a low expression of this protein in frail patients which can be justified by that lower PSRC1 levels significantly increases the levels of multiple pro-inflammatory cytokines with the activation of the NF-κB signaling pathway.

MMP-12, a human macrophage metalloelastase, is a 54kDa proenzyme which was first recognized as a protein secreted by inflammatory macrophages [27]. MMP-12 has been found to be associated with inflammatory skin disorders [28], atherosclerosis [29] and some central nervous system diseases [30]. It is reported that this protein releases TNF-α which initiate a cascade of inflammatory pathway eventually [31]. Studies suggested that MMP-12 was one of the nine circulating proteins which were associated with heart failure incidence in elderly [32]. Role of MMP-12 has also been explored in neurodegenerative disorders as it degrades myelin basic protein (MBP), a protein expressed by oligodendrocytes [33] and abnormal expression was also reported in microglial cells upon by Aβ1–42 application [34]. As reported in previous studies, MMP-12 is a pro-inflammatory mediator molecule whose abnormal expression is already reported in elderly population with risk of heart failure incidence [32] but this study first time reports the elevated levels of MMP-12 protein in frail patients which may help in creating inflammatory environment in elderly.

The process of neurogenesis decreases in presence of inflammation related diseases such as aging, neurodegenerative disorders etc. studies reported involvement of mitochondria in neurogenesis processes [35]. Mitochondria have critical role in maintaining cellular damage and leads to neurodegenerative disorders upon inactivation [36]. Mortalin is a member of HSP70 family and reported to be localized in mitochondrial matrix (MM). It is one of the most abundant protein of MM. Massa et al. [37] discovered this protein as a stress marker in central nervous system and its expression increased in relation with degree of brain injury. Other studies also confirmed its crucial role in protecting mitochondria from dysfunction and cell death [38]. Mitochondria are main source of ROS production. As mortalin is a stress marker, so upon depletion of glucose in the body, mortalin levels increase robustly to reduce the ROS accumulation which shows that mortalin can have cryoprotective effect [39]. Several studies have supported that mortalin, a stress chaperone, plays an important role by maintaining proteome integrity during aging and stressed conditions. Expression levels of mortalin were also analyzed in tissues of brain and cardiovascular diseased patients. In both cases, mortalin was found to be increased significantly with severity of diseases [37,40]. Interestingly, our results are also consistent with previous studies. In present study, increased levels of mortalin protein can be explained by considering the role of ROS production, oxidative damage, and neurodegeneration in frailty syndrome. This study may be helpful in analyzing this protein as potential biomarker for frail patients.

NK cells are necessary for providing protection to normal cells over virus-infected and cancer cells. NK cells activation is achieved by balancing signals from stimulatory receptors and MHC class I-specific inhibitory receptors [41]. Impairment in this balance will make NK cells less responsive towards infections. NKG2A are inhibitory receptors that are expressed by a majority of NK cells and on subsets of CD8+ T cells for MHC class I molecules and monitor their expression. Studies suggest that these receptors bind specifically to HLA-E ligand present on normal or infected cells which leads to the protection of infected cells against NK/T-cells mediated cell-lysis [42]. In present study, we have reported a low expression of NKG2A protein which suggests that in frailty, age-associated diseases, imbalance of inhibitory signals affect cytotoxic responses of NK cells which leads to neurodegeneration and immune system alterations [43]. Further studies need to be done in relation with frailty and this protein to confirm our results.

NME3 gene encodes Nucleoside Diphosphate kinases (NDPK) which are conserved proteins in eukaryotes. The homozygous mutation in NME3 gene was found in fatal neurodegenerative disorder [44]. The pro-inflammatory role of NDK3 in signaling downstream of TLR5, had been recognized in cancer immunotherapies [45]. Present study found 1.31 fold increase in protein levels in frail patients in comparison to non-frail patients. The role of this protein has not much explored in neurodegenerative disorders or age-associated diseases. However, this study explains that there can be a relation between this protein and frailty syndrome and it need to be explored more.

Aberrant inflammation is considered both an emerging hallmark of frailty and a supporting characteristic of ageing. Frailty is the geriatric syndrome characterized by accumulation of ROS which increases various inflammatory molecules. Some upregulated inflammatory molecules cause various types of age associated diseases whereas some molecules downregulate to prevent the body from diseases. In contrast to previous study [46]., we have found a different panel of markers in frail patients This differential expression of proteins in blood of frail elderly can serve as a significant blood based marker for the early detection of frailty and also as a target molecule for therapeutic interventions to prevent elderly from further deterioration. On the basis of present proteomics based study, we can hypothesize that frailty is associated with higher inflammatory parameters levels, in particular SAA, Haptoglobin, Mortalin, and MMP12. In order to use, these markers as potential biomarkers of frailty in the elderly, further studies are warranted.

## Conclusion

This study will help in identifying the critical pathophysiology of inflammatory pathways associated with frailty. These molecular changes can be used to identify, predict and differentiate frail from non-frail elderly. It can be concluded that this blood proteomic study of frail and non-frail elderly can aid in diagnostic assessment in clinical practice at early phase of frailty.

## Acknowledgments

Authors acknowledged all the study groups participated in this study.

